# Limits to adaptation along environmental gradients

**DOI:** 10.1101/012690

**Authors:** Jitka Polechová, Nick Barton

## Abstract

**Brief Summary:** Why do species’ ranges often end when no obvious change in the environment suggests they should? Our theory explains that there is an inherent limit to adaptation arising in any (finite) natural population, and identifies the key parameters that determine this limit to a species’ range. Two observable parameters describe the threshold when adaptation fails: i) the loss of fitness due to dispersal to a different environment, and ii) the efficacy of selection relative to stochastic effects in finite populations.

**Abstract:** Why do species not adapt to ever-wider ranges of conditions, gradually expanding their ecological niche and geographic range? Gene flow across environments has two conflicting effects: while it increases genetic variation, which is a prerequisite for adaptation, gene flow may swamp adaptation to local conditions. In 1956, Haldane proposed that when the environment varies across space, “swamping” by gene flow creates a positive feedback between low population size and maladaptation, leading to a sharp range margin. Yet, current deterministic theory shows that when variance can evolve, there is no such limit. Using simple analytical tools and simulations, we show that genetic drift can generate a sharp margin to a species’ range, by reducing genetic variance below the level needed for adaptation to spatially variable conditions. Aided by separation of ecological and evolutionary time scales, the identified effective dimensionless parameters reveal a simple threshold that predicts when adaptation at the range margin fails. Two observable parameters determine the threshold: i) the effective environmental gradient, which can be measured by the loss of fitness due to dispersal to a different environment, and ii) the efficacy of selection relative to genetic drift. The theory predicts sharp range margins even in the absence of abrupt changes in the environment. Furthermore, it implies that gradual worsening of conditions across a species’ habitat may lead to a sudden range fragmentation, when adaptation to a wide span of conditions within a single species becomes impossible.

## Introduction

Why a species’ range sometimes ends abruptly, even when the environment changes smoothly across space, has interested ecologists and evolutionary biologists for many decades [1–8]. Haldane [2] proposed that when the environment is spatially heterogeneous, a species may be unable to adapt and expand its range because gene flow from the centre swamps the populations at the range margins, preventing their adaptation. Theory showed that when genetic variance is fixed, adaptation indeed fails if the environment changes too steeply across space [9], and a sharp margin to the species’ range forms. The population remains well adapted only in the centre of the range, and gene flow swamps variants adapted to the margins, preventing range expansion.This result also elucidates range margins in the presence of competitors: then, interspecific competition in effect steepens the environmental gradient [10]. Yet, this limit to adaptation assumes that local genetic variation is fixed.

Current deterministic theory states that when genetic variance can evolve, there is no sharp limit to a species’ range [11]. The genetic mixing caused by gene flow inflates the genetic variance and facilitates further divergence. Gene flow across a phenotypic gradient maintained by the environment can generate much more variance than would be maintained by mutation alone [12, 13]. This rise of genetic variance with environmental gradient can allow species to adapt to an indefinitely wide geographic range – and hence an indefinitely wide ecological niche. Adaptation only fails when the local load due to genetic variance becomes so large that the population is no longer sustainable because the mean Malthusian fitness (growth rate) declines below zero. On constant environmental gradients, this leads to extinction everywhere. On gradually steepening gradients, we would see a gradual decrease in density due to the rising genetic variance and the associated standing load.

The interface between ecology and evolution is critical for understanding the evolution of the species’ range. Yet until now, no predictive theory has included the fundamental stochastic processes of genetic drift and demographic stochasticity. Genetic variation is often measurably lower in peripheral populations [4, 14] and experimental evidence suggests that low genetic variance coupled with high gene flow can prevent adaptation at the edge of a species’ range in nature [15–17]. Thus, there is clear evidence that low genetic variation may limit adaptive range expansion. This may be because genetic drift reduces local variance [18,19] and hence the potential of the population to adapt [20]. Studies of range expansion with genetic drift are few, and limited to simulations [21–24]. By manipulating dispersal and carrying capacity, these studies demonstrated that genetic drift may be important to theory of species’ range evolution. However, without establishing the key parameters, no quantitative predictions can be made that generalise beyond specific simulation models.

The goal of this study is to examine how finite population size, through its effect on genetic drift, and in combination with demographic stochasticity, affects the limits of a species’ range. We give the combination of parameters that describes the dynamics of evolution of a species’ range and especially, its limits, when genetic variance can evolve, the environment is heterogeneous, and populations are finite. We assume that as the environment changes across space, there is a corresponding change in the optimal value of some phenotypic trait. Deviation from the optimum reduces fitness, and so range expansion requires adaptation to this environmental gradient. Demography and evolution are considered together, and both genetic variance and trait mean can freely evolve *via* change in allele frequencies. Crucially, we include both genetic and demographic stochasticity. The model is first outlined at a population level, in terms of coupled stochastic differential equations (Materials and Methods: Evolutionary and ecological dynamics). Using this formalisation, we can identify the effective dimensionless parameters that describe the dynamics: these parameters are measurable, and predict when adaptation to an environmental gradient fails. After separation of the time-scales of ecology (fast) from evolution (slow), these reduce to just two key observable parameters – a major simplification for a model whose initial formulation requires seven parameters. Next, individual based simulations (SI: Individual-based model) determine the driving relationship between the key parameters, and test its robustness.

## Results

### Deterministic limit

First, to both validate and illustrate the model, we show that the individual-based model matches the predictions at the deterministic limit [11], where the species’ range expands indefinitely. Fig. 1 illustrates the joint evolution of population size and trait mean and variance *via* change in allele frequencies, when genetic drift is weak – when 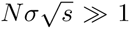 [25, 26], where population size within a dispersal distance *σ* is *Nσ*, and *s* is the strength of selection per locus. *N* represents size of each deme, which corresponds to population density in continuous space. Genetic variance, *V_G_*, evolves, and is generated primarily by gene flow due to mixing of genes from individuals with different phenotypes, well adapted to the diverse environments (Fig. 1D); the contribution from mutational variance is negligible.

**Figure 1:**
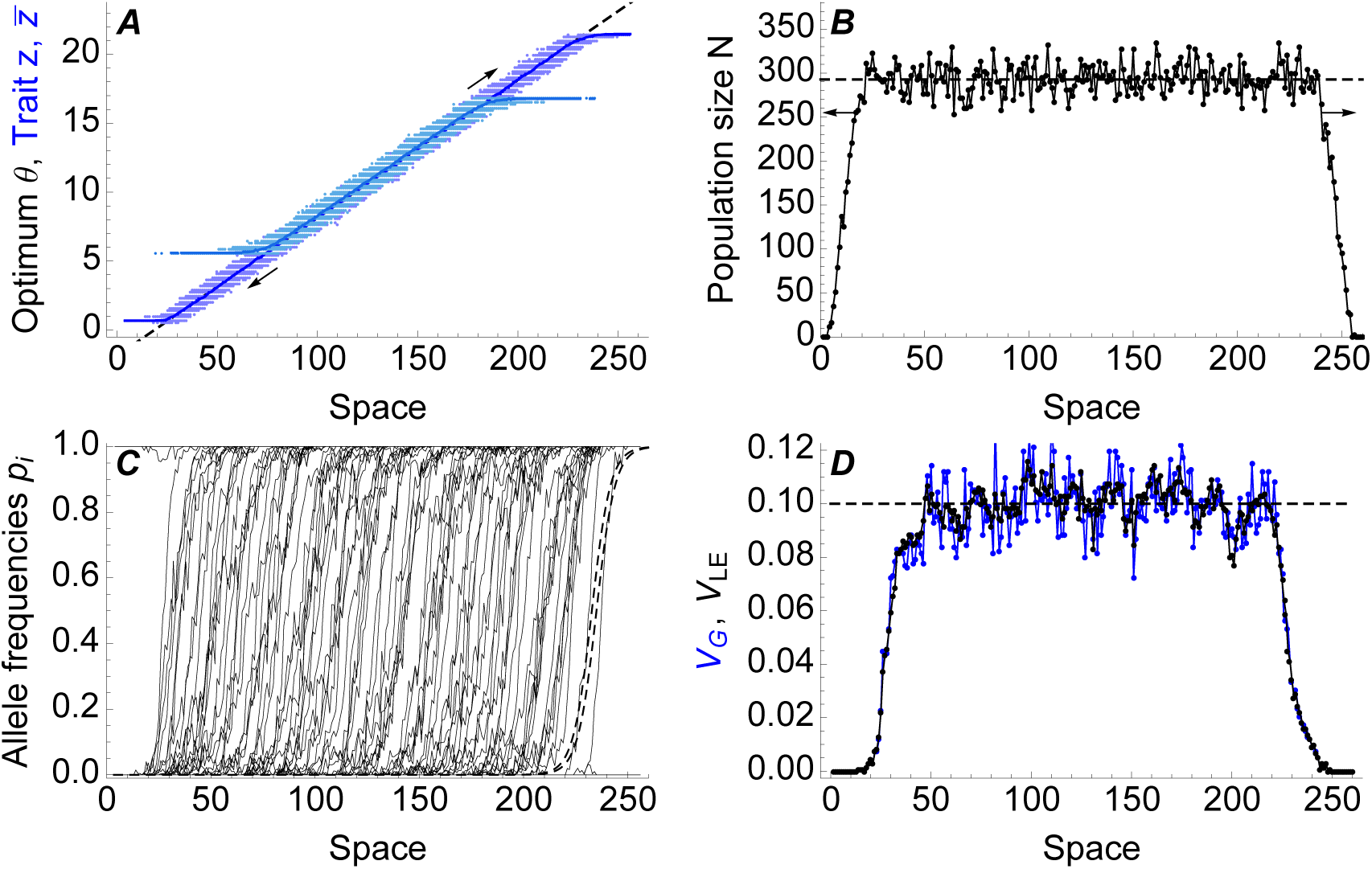
Illustration of the individual based model at the limit of weak genetic drift, when the species’ range keeps expanding as predicted by the deterministic model [11]. (A) Trait mean 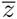 matches the optimum *θ* = *bx* (dashed line), shown for the starting population (light blue) and after 5000 generations (dark blue). The spread of the trait values *z* for all individuals is shown with dots. (B) Local population size is close to the deterministic prediction (dashed line) 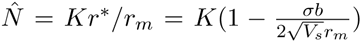, where *K* gives the carrying capacity for a well adapted phenotype. (C) Clines for allele frequencies are shown by thin black lines; the predicted clines (dashed) have widths 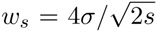 and are spaced *α*/*b* apart. (D) Total genetic variance is shown in blue, the linkage-equilibrium component in black; the dashed line gives the prediction 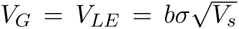: each cline contributes genetic variance 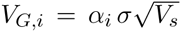 and per unit distance, there must be *b*/*α* clines if the trait mean matches into the optimum [11, p. 378]. Parameters, defined in Table 1: *b* = 0.1, *σ*^2^ = 1/2, *V_s_* = 2, *r_m_* = 1.025, *K* = 300, 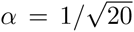, *μ* = 10^−6^, 5000 generations.

**Table 1:**
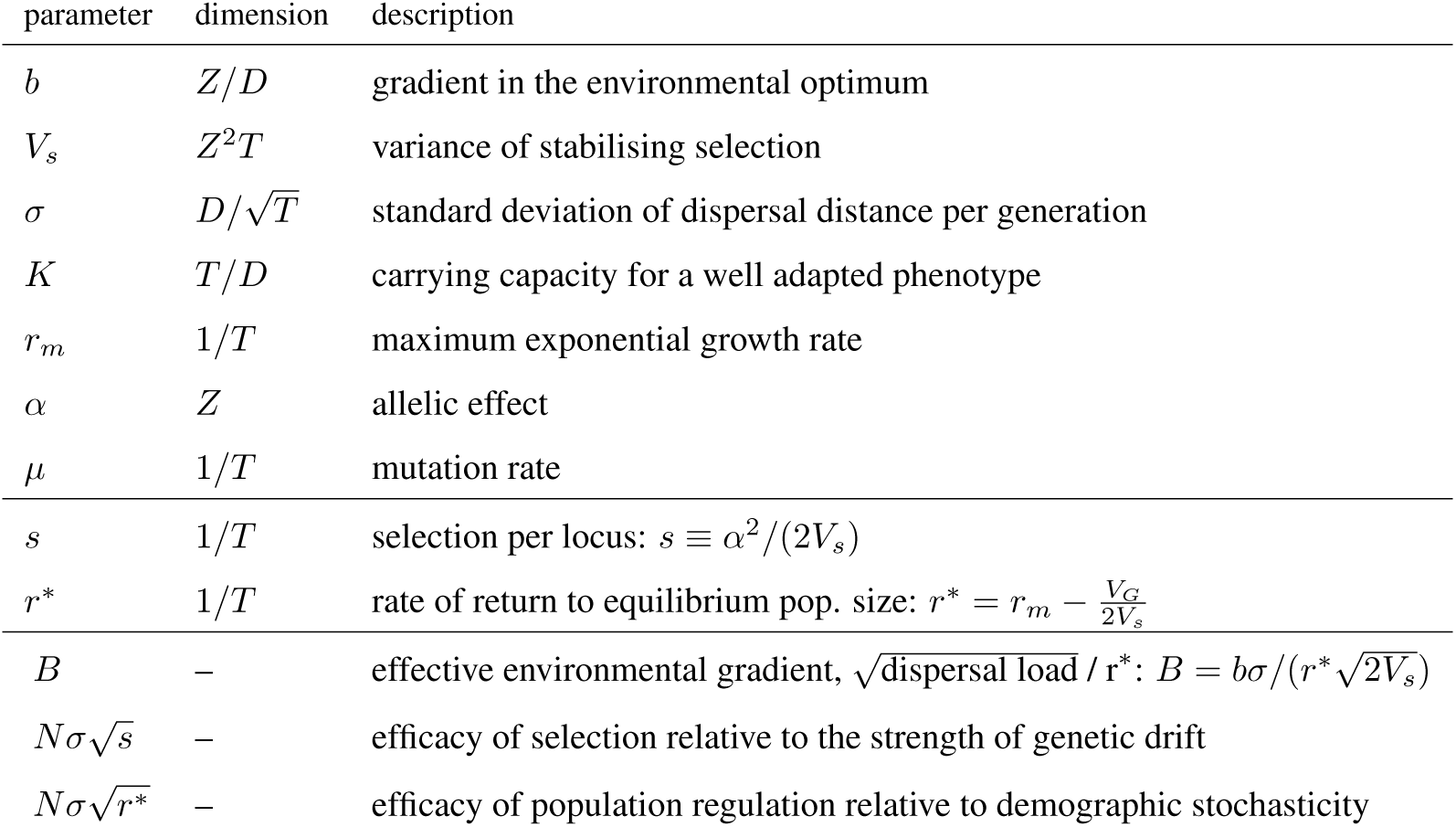
Three scale-free parameters: *B*, 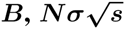 and 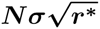 describe the system. Top section gives seven parameters of the model before rescaling, middle section gives important composite parameters. Denoting the dimensions, T stands for time, D for distance and Z for trait. Note that with a Poisson number of offspring, the effective population size *N_e_* (which measures rate of genetic drift / coalescence) is identical to the *N* that regulates population growth due to crowding: hence both carrying capacity *K* and population size *N* have units of *T*/*D*. Mutation rate *μ* is set to be small, with minimal contribution to the dynamics, and hence *μ*/*r*^*^ is neglected in the rescaled parameterization (bottom).

### Scaling and separation of time scales

We now proceed by reducing the number of parameters (Table 1) in the model, including stochasticity (Materials and Methods: Rescaling). We seek a set of dimensionless parameters that fully defines the dynamics of a system, with a preference for those that are empirically measurable. Rescaling space, time, trait, and population density reveals that three dimensionless parameters fully describe the system, neglecting mutation and assuming linkage equilibrium between loci. The first parameter carries over from the phenotypic model of [9]: i) the effective environmental gradient, 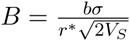, where *b* is the gradient *V_S_* in the optimum, *V_s_* the width of stabilising selection, and *r*\* gives the strength of density-dependence. Two additional dimensionless parameters come from including stochasticity: ii) the efficacy of population regulation relative to demographic stochasticity, 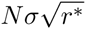; and iii) the efficacy of selection relative to the strength of genetic drift, 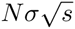.

The dynamics of evolution of species’ range simplifies further because selection per locus *s* is typically much smaller than the rate of return to equilibrium population density *r*\* (see [27] and [28, Appendix D]; Fig. S1 for values used here). This has two consequences. First, the ecological dynamics *∂N*/*∂t* are faster than the evolutionary dynamics *∂p*/*∂t* of the individual loci (where *p* denotes the allele frequency and *t* denotes time). Importantly, the genetic variance evolves slower than the mean: whereas the trait mean changes with the product of strength of selection and the genetic variance, the directional change in the variance is slower by the inverse of the number of locally polymorphic loci (precisely, the “effective number of loci” [29]). Genetic drift only slowly degrades expected heterozygosity 〈*pq*〉 at each locus (Fig. S2), reducing the variance. Hence, over ecological time scales, genetic variance can be treated as constant. Second, the effect of fluctuations due to genetic drift, scaling in one-dimensional habitat with 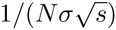 (see Fig. S2), is expected to dominate over the effect of demographic fluctuations, that rise with 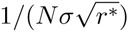. Hence, we expect that the conditions determining whether adaptation to an environmental gradient is sustainable within a single species’ will be described by just two of the dimensionless parameters. These are the effective environmental gradient *B* and the efficacy of selection relative to the strength of genetic drift 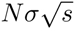. We thus propose that the dynamics of species’ range evolution can be understood based on bulk parameters *B* and 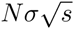, rather then by focusing on asymmetric gene flow near the range margins. Later, we show that the prediction holds even when *B* and 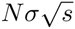 change steadily across space.

### Threshold for collapse of adaptation

First, we simulated the basic model with a linear gradient, assuming equal phenotypic effects *α* of each allele, and including both genetic and demographic stochasticity. Parameters were drawn at random from distributions consistent with our knowledge of the range of parameters expected in nature [28, Discussion] and ensuring that without genetic drift, all ranges would expand (see Fig. S1). Then, we performed additional runs to test whether the threshold obtained from the linear gradient applies when parameters are changing gradually across space. Namely, we tested whether a stable range margin forms at the predicted value when the environmental gradient varies across space or when the carrying capacity is non-uniform. Last, we tested the model assuming that allelic effects *α_i_* are exponentially distributed.

Fig. 2 shows our key result: the effective environmental gradient, *B*, and the efficacy of selection relative to the strength of genetic drift, 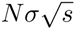, determine the threshold for collapse of adaptation. This is because the genetic variance evolves primarily in response to 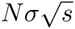 and *B*, whilst the effect of demographic stochasticity, 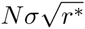, is relatively weak. When 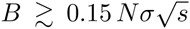, genetic drift strongly degrades adaptation to a steeply changing environmental optimum, and the species’ range contracts. The constant 0.15 is obtained as the best fitting threshold for the data in Fig. 2. At the start of all simulations, the population is perfectly adapted in the central part of the available habitat. As both environmental gradient and carrying capacity are uniform across the habitat, populations either steadily expand or contract. Typically, populations contract from their margins and ultimately (over times much longer than 5000 generations of the simulation runs) collapse to a state with no or very little clinal variation. However, when genetic drift is very strong, populations collapse abruptly or even fragment (Fig. S3): this second threshold, based on a “critical gradient” of a phenotypic model, is explained later. This model with an idealised linear gradient and uniform carrying capacity is used to identify the relationship between the driving parameters 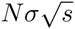 and *B*, which determine whether a population can adapt to an environmental gradient.

**Figure 2:**
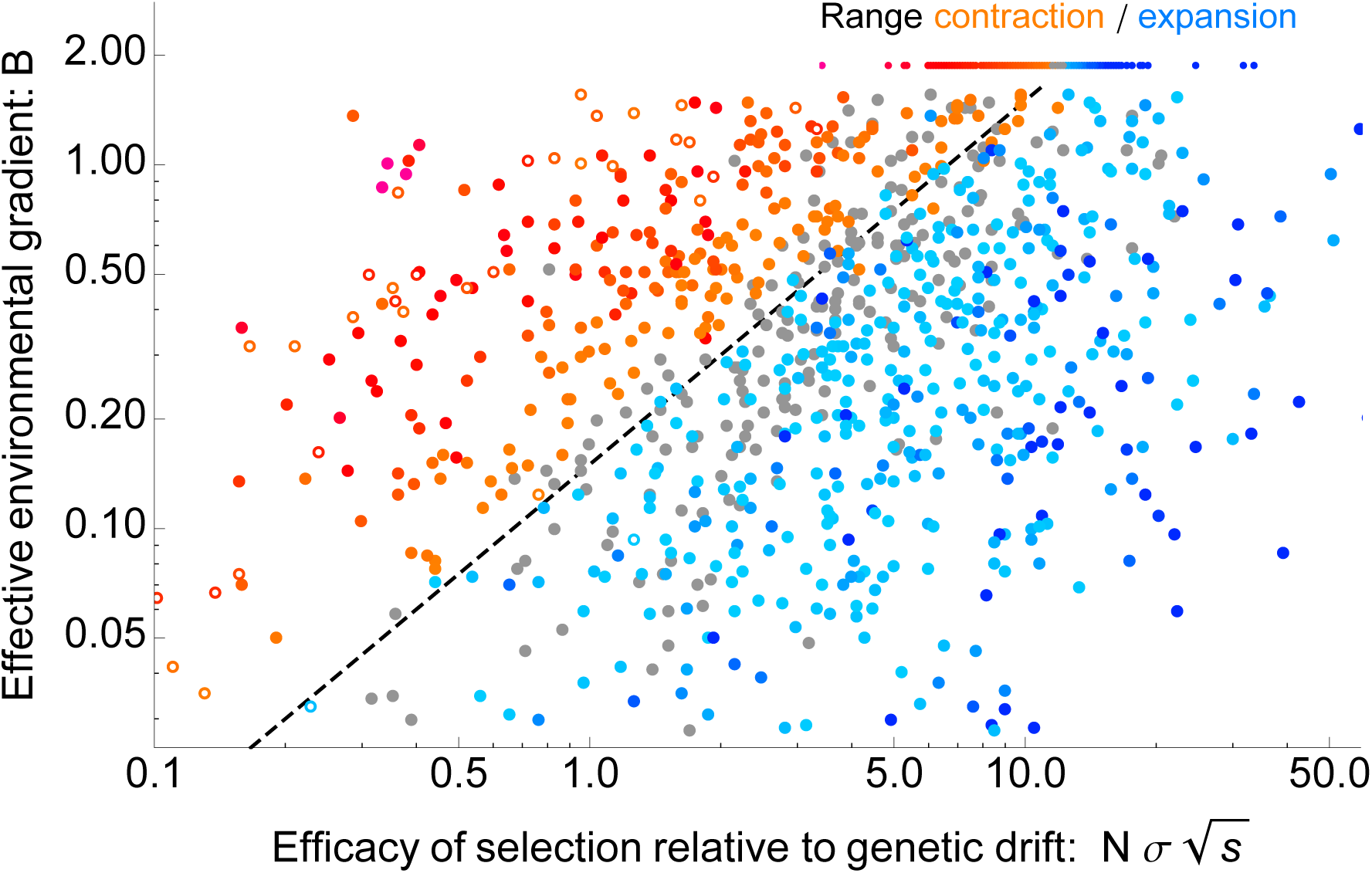
Species’ range starts to contract when the effective environmental gradient is steep compared with the efficacy of selection relative to genetic drift: 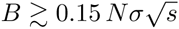, where 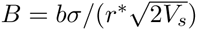. This threshold is shown by a dashed line. The rate of expansion increases from light to dark blue and rate of range contraction increases from orange to red. Grey dots denote populations for which neither expansion nor collapse is significant at *α* = 2%. Open dots indicate fragmented species’ ranges (illustrated by Fig. S3). The ranges of the underlying (unscaled) parameters are in the following intervals: *b* = [0.01, 1.99], *σ* = [0.5, 4.8], *V_s_* = [0.006, 8.4], *K* = [4, 185], *r_m_* = [0.27, 2] and *α* = [0.01, 0.39], *μ* = [10^−8^, 8 · 10^−5^]; the number of genes is between 7 and 3971. The selection coefficient per locus is hence in the interval of *s* = [3 · 10^−4^, 0.66], with median of 0.007. Parameter distributions are shown in Fig. S1.

In reality, the environment does not vary precisely linearly, and the carrying capacity is not uniform. When the steepness of the environmental gradient varies steadily across space, this threshold, 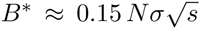, indicates where a stable range margin forms (Fig. 3). Without genetic drift, the genetic variance would steadily inflate with increasing environmental gradient, gradually reducing local population size due to an increasing number of maladapted individuals (see dashed lines in Fig. 3D). With genetic drift, the variance is pushed below the level necessary to maintain adaptation, the trait mean abruptly fails to match the optimum, and so a sharp margin to the range forms. Similarly, a sharp range margin forms when 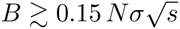, if carrying capacity declines across the habitat for extrinsic reasons (Fig. S4). Species’ range is more robust to spatial fluctuations in carrying capacity within the occupied habitat. Whilst range expansion stops at the predicted threshold, a (locally) large fall in density below the threshold is necessary for the range to fragment within a previously occupied habitat: see Fig. S5.

**Figure 3:**
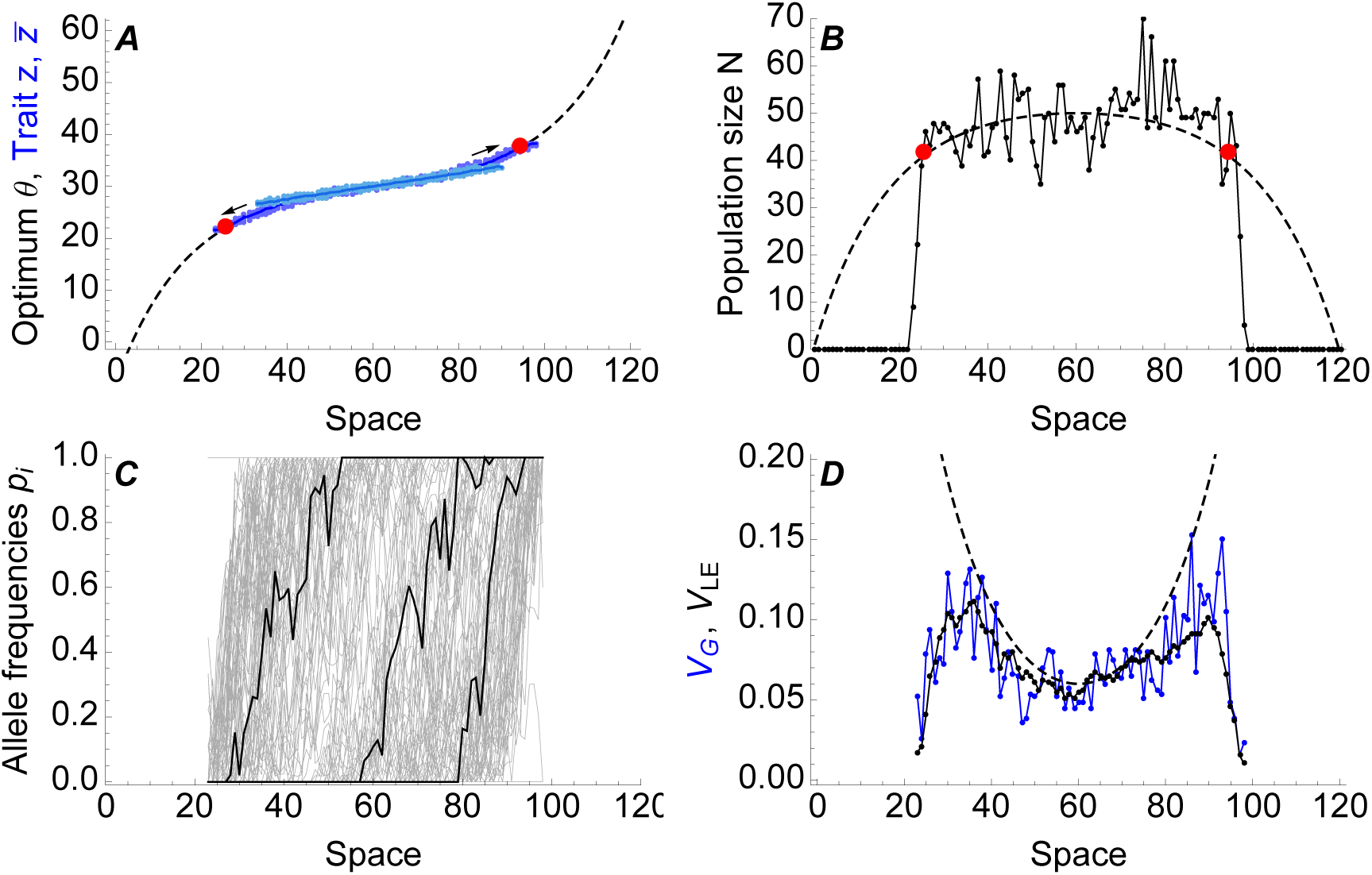
With a steepening environmental gradient, a stable range margin forms when 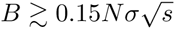 (red dots). (A) The gradient in trait mean follows the environmental optimum (dashed line) until the gradients steepens so that 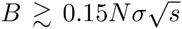, where expansion stops. (B) Population density drops off sharply when the predicted threshold (red dots) is reached. Dashed line gives the predicted population size assuming variation is not eroded by genetic drift. (C) Three representative clines are shown in black, other clines form the gray background. (D) Adaptation fails when genetic variance fails to increase fast enough to match the steepening environmental gradient (total variance *V_G_* in black, linkage-equilibrium component *V_LE_* in blue). For all subfigures, dashed lines give deterministic predictions ( [11] and Fig. 1). Parameters: central gradient *b*_0_ = 0.12, *σ*^2^ = 0.5, *V_s_* = 0.5, *r_m_* = 1.06, *K* = 53, *μ* = 2 · 10^−7^. Time = 100 000 generations; expansion stops after 40 000 generations; (A) also shows the initial stage in light blue.

A sharp range margin forms not only when all loci have equal allelic effects (i.e., the trait changes by a fixed value due to every substitution), but also when allelic effects are exponentially distributed. Then, range expansion slows down progressively around the threshold (Figs. S6 and S7), described by 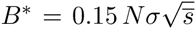, where the mean selection coefficient is 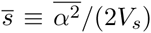. The mean selection coefficient gives an estimate for the expected range margin because clines at weakly selected loci are degraded by genetic drift (Fig. S2 and [26]), reducing the genetic variance. For the population to expand further beyond the threshold, positively selected alleles with increasingly large effect need to arise (Fig. S7; see also [30]). In natural populations, these become ever rarer; and for any finite distribution of allelic effects (as in our model), are exhausted.

In the absence of genetic drift, low dispersal can enhance adaptation by reducing swamping by gene flow [11,16]. However, with genetic drift, this is no longer true. To a first approximation, both the efficacy of selection relative to genetic drift 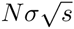 and the effective environmental gradient 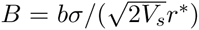 increase at the same rate with dispersal. Only a weak dependence on dispersal remains via *r*\* (because 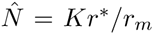 and *r*\* decreases with increased genetic load due to mixing across the gradient), which favours low to intermediate dispersal: see Fig. S9; *cf.* [23]. The beneficial effect of intermediate dispersal can be stronger in one-locus models, because in the absence of recombination, dispersal only increases crowding in the new habitat [31] and flow of the advantageous mutants back to the old habitat [32] whilst bringing new targets for mutation and positively selected mutants. Increased dispersal can thus enhance fitness at the range edge even in the absence of inbreeding depression – *cf.* [33].

A second threshold, above which the population collapses abruptly, can be understood from a deterministic phenotypic model with fixed genetic variance [9]. This theory showed that when genetic variance *V_G_* is fixed, there is a critical environmental gradient 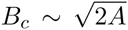 (where *A* = *V_G_/*(*r*\**V_s_*)), above which the trait mean fails to track the spatially changing optimum, and the population is well adapted only in the centre of its range (see [9, p.6] and [28, Fig. 3]). However, such a limit does not exist when genetic variance evolves due to gene flow across environments [11] in the absence of genetic drift. We show that genetic drift can reduce the variance such that the “critical gradient” *B_c_* is reached, despite the variance evolving: then, population abruptly loses most of its genetic variation and suffers a demographic collapse (Fig. 4). This demographic collapse may lead to either a rapid extinction from most of the species’ range or to range fragmentation, when a species cannot maintain adaptation throughout the whole range, yet multiple isolated populations persist (Fig. S3).

**Figure 4:**
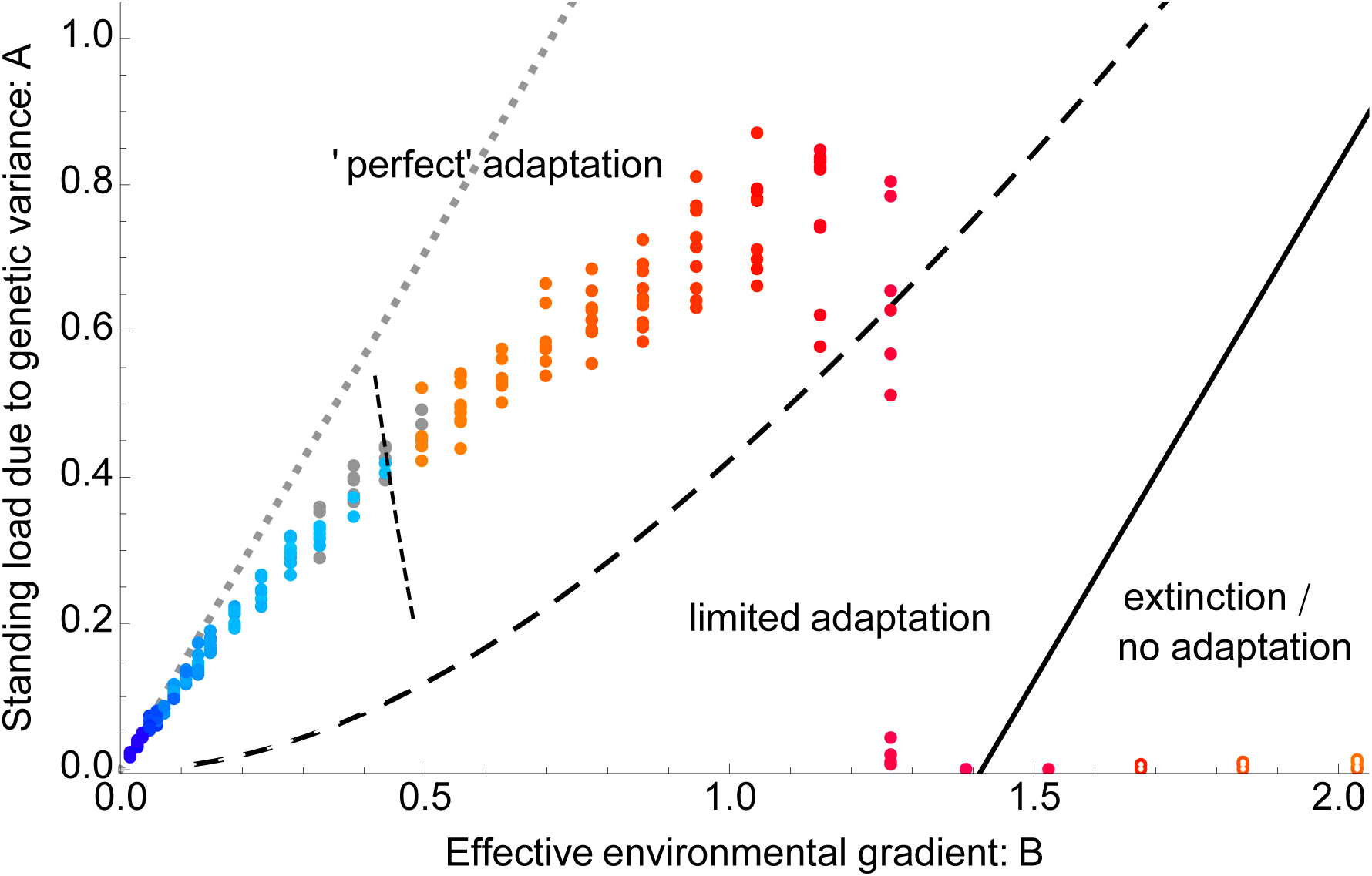
The phenotypic model predicts a second sharp transition in the dynamics (*B_c_*, dashed curve). As the effective environmental gradient 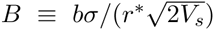 increases, the scaled variance *A* ≡ *V_G_*/(*r*^*^*V_s_*) increasingly deviates from the deterministic prediction with evolvable variance ( [11], gray dotted line). The variance decreases due to the combined forces of genetic drift (see Fig. S2 A) and selection on small transient deviations of the trait mean from the optimum. Once 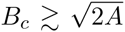 (dashed curve), the population collapses abruptly. Furthermore, no adaptation is maintained beyond the solid line 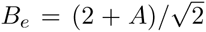, where the phenotypic model [9] predicts extinction. Open dots (lower right) denote fragmented species’ range (Fig S3). The colours are as in Fig. 2, the threshold when range starts to contract, 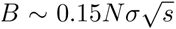, is illustrated by the short steep dashed line. Parameters: *b* increases from 0.025 to 1.25, *σ*^2^ = 1/2, *V_s_* = 1/2, *K* = 50, *α* = 1/10, *r_m_* = 1.06, 5000 generations. Ten replicates are shown for each *B*.

## Discussion

We are just becoming to be able to assess the genomics of local adaptation across a species’ range [34]. Hence, there is a great need for a theory that integrates both genomic and demographic data. It would be exciting to see how our prediction improves experimental tests of the causes of range limits in one-dimensional habitats such as along a deep valley or a river, such as [16], that assumed fixed genetic variance [9]. The predictive parameters can in principle be measured: individuals exactly one standard deviation for the distribution of parent-offspring dispersal distances along the environmental gradient gives a loss of fitness *B*^2^/*r*^*^. The parameter *r*\*, which describes the rate of return of population to equilibrium can be estimated (eg., following [35]). Neigbourhood size in a one-dimensional habitat, 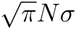, can be estimated from neutral markers [36, Eq. 2]. The efficacy of selection relative to genetic drift, 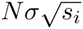, will vary across loci. The selection coefficients *s_i_* = *α_i_*^2^/(2*V_s_*) can then be estimated either by mapping allelic effects *α_i_* from the QTLs (quantitative trait loci) underlying an adaptive trait and measuring the strength of stabilising selection 1/(2*V_s_*), or by estimating the selection per locus directly from the steepness of the clines in allele frequencies across space. Steepness of clines gives the desired *s_i_* even under pleiotropy, whereas the first measure gives the stabilising selection due to all pleiotropic effects. Given a fixed (finite) distribution of selection coefficients *s_i_*, we can predict when adaptation is expected to fail: for example, when, given a reduction in population size, a species’ range would become prone to fragmentation.

Our main result, that adaptation fails when the effective environmental gradient is large relative to the efficacy of selection versus genetic drift, 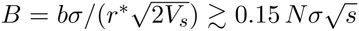, can be rephrased to a form that is closely related to Haldane’s cost of selection [37]. Haldane showed that in a single population, each substitution requires a certain number of selective deaths (i.e., reduction in mean fitness relative to the maximum possible), which is nearly independent of the strength of selection per locus. In our model with a spatially varying environment, *b*/*α* substitutions are required per unit distance: as we assume hard selection, these substitutions need to also arise via selective deaths. If too many selected substitutions are needed relative to births in the population, such that *b*/*α* ≳ 0.15 *N r*\*, adaptation fails (Fig. S8). However, this failure is due to stochastic fluctuations, and so depends on the effective number of deaths per generation, *Nr*\*. Note that a sharp range margin cannot form when selection is soft – when *K* fittest phenotypes are always selected, a population spreads as long as its fitness is above zero.

This intrinsic limit to adaptation was found by analysing one-dimensional habitats: whilst initial simulations show that all driving parameters are preserved for a narrow two-dimensional habitat, as habitats become wide (≳ 100 *σ*), the effect of genetic drift changes: stochastic fluctuations of allelic frequencies become only weakly dependent on selection [38]. The range limits for broad two-dimensional habitats will be the subject of a future paper.

Our theory shows that there is an inherent limit to adaptation arising in any (finite) natural population, and identifies the key parameters that determine this limit to a species’ range. It explains that a sharp range margin forms when fitness cost, induced by a spatially varying environment, becomes too high relative to the efficacy of selection in the presence of genetic drift – even in the absence of fixed genetic constraints, such as insufficient genetic variance [9, 39] or rigid fitness trade-offs between traits [40, 41]. Because the threshold depends only on the fitness cost of dispersal and the efficacy of selection per locus relative to genetic drift, it readily generalises to many traits. It gives an upper limit: even in the absence of trade-offs (whether transient or rigid), adaptation fails at the estimated threshold. Throughout, we have assumed additivity between the allelic effects: that is consistent with the experimental evidence, suggesting that statistical epistasis is typically weak [42].

Within an already occupied habitat, small fluctuations in carrying capacity below the predicted threshold do not lead to range fragmentation. Nevertheless, the coherence of the species’ range is sensitive to habitat fragmentation: a local but large fall in carrying capacity facilitates extinction from its boundaries wherever the threshold has been crossed (Fig. S5). Swamping by gene flow at the existing margins, as emphasised by [2], is important for the formation of a sharp range edge – even though the local bulk parameters *B* and 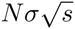 predict where the range margin forms. Our theory is entirely consistent with the lack of evidence for an abundant centre [43] and with only a small decrease in neutral diversity in peripheral populations [14]: we show that even a small decrease in the attainable equilibrium density (carrying capacity) can lead to a sharp range margin.

The theory also implies that gradual decrease in carrying capacity can lead to a collapse of adaptation as genetic drift erodes the genetic variation, causing a sudden collapse or a fragmentation of a species’ range. When the effective environmental gradient *B* varies across space, steeper sections would act as barriers to a species’ spread – in contrast to the deterministic model [11]. This is important for management of biological invasions when adaptation is required [44, 45] and for biological conservation [46]: a species’ range may collapse due to genetic drift well before demographic factors (emphasised in [47]) become significant [46]. Understanding species’ range limits in a constant environment is also essential before extending the model to account for temporally changing environments, such as when modelling joint adaptation and range expansion in species’ responding to climate change [48]. Furthermore, the predicted emergence of inherent limits to species’ ranges across steadily varying environments offers extensions to the theory of ecological speciation, and eventually, may help us to elucidate macroecological patterns of biodiversity.

## Materials and Methods

### Evolutionary and ecological dynamics

We model the joint evolution of i) population size and ii) trait mean and its variance *via* change in allele frequencies. The fitness of an individual declines quadratically with the deviation of the trait *z* from an optimum *θ* that changes linearly across space: *θ* = *bx*, where *b* is the gradient in the environment and *x* is the distance in one dimension. The phenotypic trait *z* is determined by many additive di-allelic loci, so that genetic variance can evolve. For simplicity, we use a haploid model: for additive allelic effects, the extension to a diploid model is straightforward.

The Malthusian fitness of a phenotype *z* is *r*(*z*, *N*) = *r_e_*(*N*) + *r_g_*(*z*), where *r_e_*(*N*) is the growth rate of a perfectly adapted phenotype, and includes density dependence; *r_g_*(*z*) ≤ 0 is the reduction in growth rate due to deviation from the optimum. *N* is the population density. The ecological component of growth rate *r_e_* can take various forms: we assume that regulation is logistic, so that fitness declines linearly with density *N*: *r_e_* = *r_m_*(1 − *N*/*K*), where *r_m_* is the maximum per capita growth rate in the limit of the local population density *N* → 0. The carrying capacity *K* (for a perfectly adapted phenotype) is assumed uniform across space. Stabilising selection on the optimum *θ* has strength 1/(2*V_s_*). Hence, for any individual, the drop in fitness due to maladaptation is *r_g_*(*z*) = −(*z* − *θ*)^2^(2*V_s_*). The genetic component of the local mean fitness is then 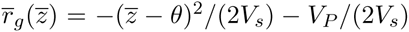, where *V_P_* = *V_G_* + *V_E_* is the phenotypic variance. The loss of fitness due to environmental variance *V_E_* can be included in 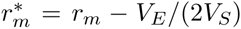; hence in this model, *V_E_* is a redundant parameter. We assume that selection is hard: the mean fitness – and hence attainable equilibrium density – decreases with increasing maladaptation: 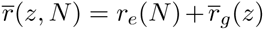. I.e., we assume that resources become more difficult to acquire and process as maladaptation increases.

For any given additive genetic variance *V_G_* (assuming a Gaussian distribution of breeding values), the trait mean 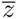 satisfies:

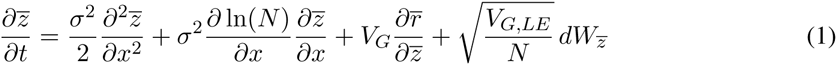

The first term gives the change in the trait mean due to migration with mean displacement of *σ*, the second term describes the effect of the asymmetric flow from areas of higher density and the third term gives the change due to selection [49, Eq. 2]. Following [25], and using that genetic variance at linkage equilibrium is 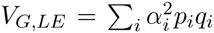. The last term gives the fluctuations in the trait due to genetic drift over infinitesimally short times *dt*: *dW*_*_ represents white noise, uncorrelated in space and time, with expectation 〈*dW*_*_〉 = 0 and 〈*dW*_*_(*x*, *t*) *dW*_*_(*x*′, *t*′)〉 = *δ*(*x*−*x*′)*δ*(*t*−*t*′)*dt dx*; *δ* is the Dirac delta [50].

Assuming additivity between loci, the trait mean is 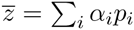 for a haploid model, where *p_i_* is the *i*-th allele frequency, *q_i_* = 1 − *p_i_* and *α_i_* is the effect of the allele on the trait – the change of the trait mean 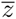 as frequency of locus *i* changes from 0 to 1. (A diploid model would have 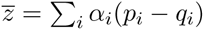.) The equation for the change of allele frequencies *p_i_* is the same for both haploid and diploid models:

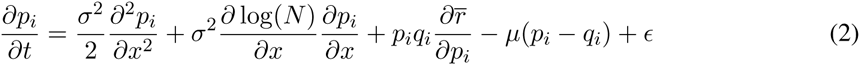

The expected change of allele frequency due to a gradient in fitness and local heterozygosity is 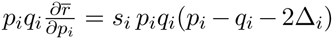, where selection at locus *i* is 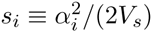 and 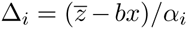 [11, two-allele model, Appendix 3]. The fourth term describes the change due to (symmetric) mutation at rate *μ*. The last term 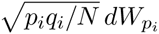 describes fluctuations in allele frequencies due to genetic drift (following [25]). Equation 2 is only exact at linkage equilibrium (i.e., neglecting covariance between allele frequencies). This is a good approximation for unlinked loci: whereas migration across the habitat generates positive linkage disequilibrium between any pair of loci, stabilising selection drives negative disequilibrium, and these cancel precisely unless selection per locus is strong. The derivation, which generalizes an ingenious but little known argument by Felsenstein [51], is given in SI: Linkage equilibrium.

Population dynamics reflect diffusive migration, growth due to the mean Malthusian fitness 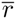, and stochastic fluctuations. The number of offspring follows a Poisson distribution with mean and variance of *N*. Fluctuations in population numbers are given by 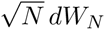 (e.g. [52]), where *dW_N_* describes an independent white noise:

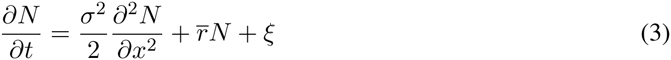

### Rescaling

The model can be further simplified by rescaling time *t* relative to the rate of return to equilibrium population size, *r*\* (which we consider equivalent to the strength of density-dependence; see also [35]), distance *x* relative to dispersal *σ*, trait *z* relative to strength of stabilising selection 1/(2*V_s_*) and local population size *N* relative to equilibrium population size with perfect adaptation 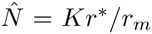; as in [9, 11]. The scaled dimensionless variables are *T*= *r*\**t*, 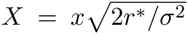, 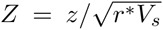 and 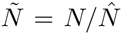. The rescaled equations for evolution of allele frequencies and for demographic dynamics then are:

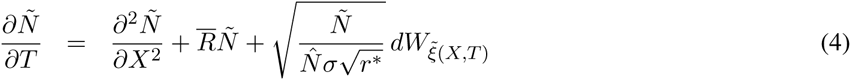

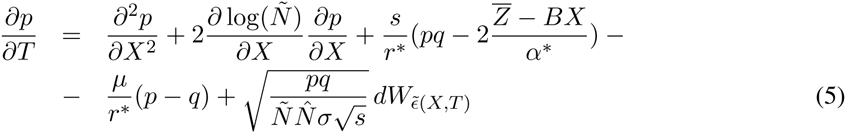

where 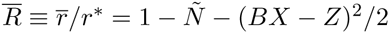 and 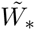 describes the independent Wiener processes in scaled time *T* and space *X*.

The rescaled equations 4 and 5 show that four parameters fully describe the system. The first two are i) the effective environmental gradient, 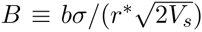 and ii) the strength of genetic drift relative to selection, near the equilibrium 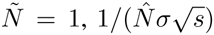. As a third parameter, one can either take iii) demographic stochasticity relative to strength of density-dependence 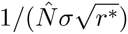 or the ratio of 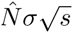 to 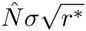, which is the strength of selection relative to density dependence, *s*/*r*\*. (The scaled effect of a single substitution, 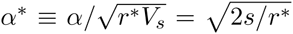.) The effect of this third parameter is small, because *s* ⋘ *r*\*. The fourth parameter, *μ*/*r*^*^, will typically be very small, and hence we will neglect it. Note that 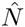 gives population density, where population is still well adapted: 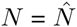 throughout most of the species’ range away from sink regions near the existing margins. Table 1 (bottom) summarises the full set that describes the system.

## Acknowledgments

We would like to thank S. Baird, A. Betancourt, J.P. Bollback, J.R. Bridle, D. Field, A. Hancock, F. Jesse, T. Paixão, T. Priklopil, R.A. Nichols, S. Novak, S. Sarikas, J.R.G. Turner, H. Uecker and M.G.J. de Vos for discussions and comments on the earlier drafts, European Research Council grant (250152) to N.B. for funding and the editor and the referees for their valuable comments.

## SI: Individual-based model

We test the predictive power of the derived key parameters using individual based simulations. Individuals are distributed among demes that form a one-dimensional habitat, with the phenotypic optimum varying along the habitat, and experience a life cycle consisting of selection, mutation, recombination, and then dispersal. Distance is measured in demes, time in generations. Every generation, each individual mates with a partner drawn from the same deme, with probability proportional to its fitness, to produce a number of offspring drawn from a Poisson distribution with mean of *Exp*[*r*], where *r* is the individual’s Malthusian fitness in continuous time. Generations are non-overlapping. Individual fitness declines due to deviation of the phenotypic trait (*z*) from the optimum and due to crowding (fitness is density-dependent). The trait is determined by a number of additive di-allelic loci, which permits genetic variation to evolve. All parameters are described in Table 1. The model is derived as a limit to continuous time, and so applies to a wide range of models that reduce to this limit. In this limit, the rate of spatial dispersal depends only on the variance of distance moved, and the effective population density (for both allele frequency and demographic fluctuations) depends only on the variance of offspring number.

**Life cycle:** Discrete-time individual based simulations are set to correspond to the model with continuous time and space. The life-cycle is selection → mutation → recombination → migration.

**Dispersal:** The habitat is formed by a one-dimensional array of demes. With deme spacing *δx* = 1, the population size per deme corresponds to the population density. We assume diffusive migration with a Gaussian dispersal kernel. The tails of the dispersal kernel need to be truncated: we choose truncation at two standard deviations of the dispersal kernel throughout, and adjust the dispersal probabilities following [1, p. 1209] so that the discretised dispersal kernel sums to 1, and the variance of dispersal is adjusted correctly. For dispersal per generation at 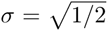, dispersal reduces to a nearest neighbour migration with a probability of migration left and right of *m* = 1/4.

**Selection:** Every generation, each individual produces a Poisson number of offspring with mean of the individual’s fitness *Exp*[(*r*)]; where 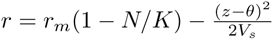, as defined earlier.

**Mutation:** Mutation rate is set to be small so that its contribution to genetic variance is negligible, but large enough to in principle enable expansion of a species’ range over the total time of 5000 generations. Specifically, it is set to one substitution per the whole population and generation. Genetic variance maintained in a population due to dispersal across environments can be substantially larger than genetic variance maintained by mutation-selection balance in uniform environments. In uniform environments, mutational variance 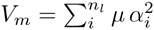 (where *α_i_* are the allelic effects, and *n_l_* is the number of loci) is robustly estimated to be between about 5 · 10^−5^*V_E_* and 5 · 10^−3^*V_E_* [2]. Taking a heritability *h*^2^ ≡ *V_G_*/(*V_G_* + *V_E_*) = 1/3 we get *V_m_* between 10^−4^*V_G_* and 10^−2^*V_G_*. In our model, genetic variance is inflated due to dispersal across environments, and so *V_m_/V_G_* must be yet smaller. Taking the higher limit of *V_m_* = 10^−2^*V_G_*, follows that *μ* should be smaller than about 4 · 10^−3^. It turns out that in general, the increase of genetic variance due to mutation cannot be fully included in the predictions, as the contribution of mutation–selection balance cannot be robustly separated from the clinal variation. A considerably higher genetic variance than *V_G_*_,*mut*_ = 2*µ n_l_ V_s_* (up to the limit of 1/4*α*^2^*n_l_*) can arise due inflation of the variance by mutation along existing clines. Therefore, we concentrate on a parameter range where the contribution of mutation to genetic variance is low, which is a biologically plausible range. In our model, genetic variance is maintained by gene flow across the environment.

**Reproduction, recombination:** The mating partner is drawn from the same deme, with the probability proportional to its fitness. Selfing is allowed at no cost. The genome is haploid with unlinked loci (the probability of recombination between any two loci is 1/2); the allelic effects *α_i_* of the loci combine in an additive fashion.

**Simulation runs:** Evolution starts with a well adapted population at the centre of the habitat. The habitat is about 10 cline widths wide; the number of genes is chosen so that with all genes adapted, the population spans the whole habitat, and that there are enough genes to maintain the “optimal” variance 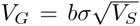 at the central part of the habitat. At the start of the simulation, half of the genes are adapted: their clines take the form and spacing as assumed for the deterministic model under linkage equilibrium.

The population evolves for 5000 generations; in total, we recorded over a thousand runs where without genetic drift the local population density would be greater than 4 assuming uniform adaptation (such that trait mean matches the optimum). We a-priori eliminated very small local population sizes (*N* < 4) so that the population size within a generational dispersal is not excessively small. The parameters were first varied one at a time, and then we tested the threshold drawing the parameters from distributions consistent with our knowledge of the range expected in nature (see [3, Discussion]). The latter will be referred to as the “random” set, with 1000 runs. The Mathematica code for the simulations, including the distributions used to draw the unscaled parameters for the “random” set is provided in FileS1. Fig. S1 shows the realised distributions for both the unscaled parameters and compound parameters.

## SI: Linkage equilibrium

Stabilising selection on a quantitative trait generates negative linkage disequilibrium, whereas dispersal generates positive linkage disequilibrium. Felsenstein’s [4] analysis of variance components showed that at equilibrium, the linkage disequilibrium generated by dispersal cancels out with the negative linkage disequilibrium generated by stabilising selection.

The argument extends to a quantitative trait determined by many di-allelic loci, here demonstrated for a haploid genome. The genetic variance is 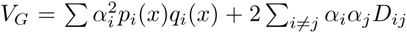. The increase of linkage disequilibrium at QLE [5, quasi-linkage equilibrium] between dispersal with variance *σ*^2^ and recombination *r_ij_* is given by 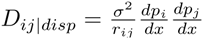 [7]. With allele frequencies at equilibrium, the linkage disequilibrium generated by stabilising selection alone is 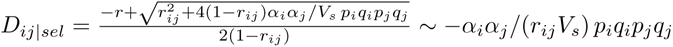 for *D_ij_* small.

In this first order approximation, the terms cancel for each pair of loci when the cline shape is the same as that under linkage equilibrium [6, two-allele model] – independently of the cline spacing across space: *D_ij_* ≡ *D_ij_*_|*disp*_ + *D_ij_*_|*sel*_ = 0. This is because 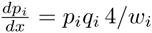, and the cline width at linkage equilibrium is 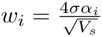.

It may be that the cline width *w_i_* or the linkage disequilibrium is distorted by additional forces, and/or by strong selection – then the linkage disequilibrium components due to dispersal and selection may not cancel. However, unless selection is strong, the first order approximation gives a simple prediction for the pairwise disequilibrium: 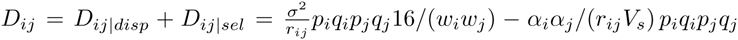. For example, we can expect that positive linkage disequilibrium *D_ij_*, generated by long-range dispersal, would drive steeper clines.

## SUPPLEMENTARY FIGURES

**Figure S1:**
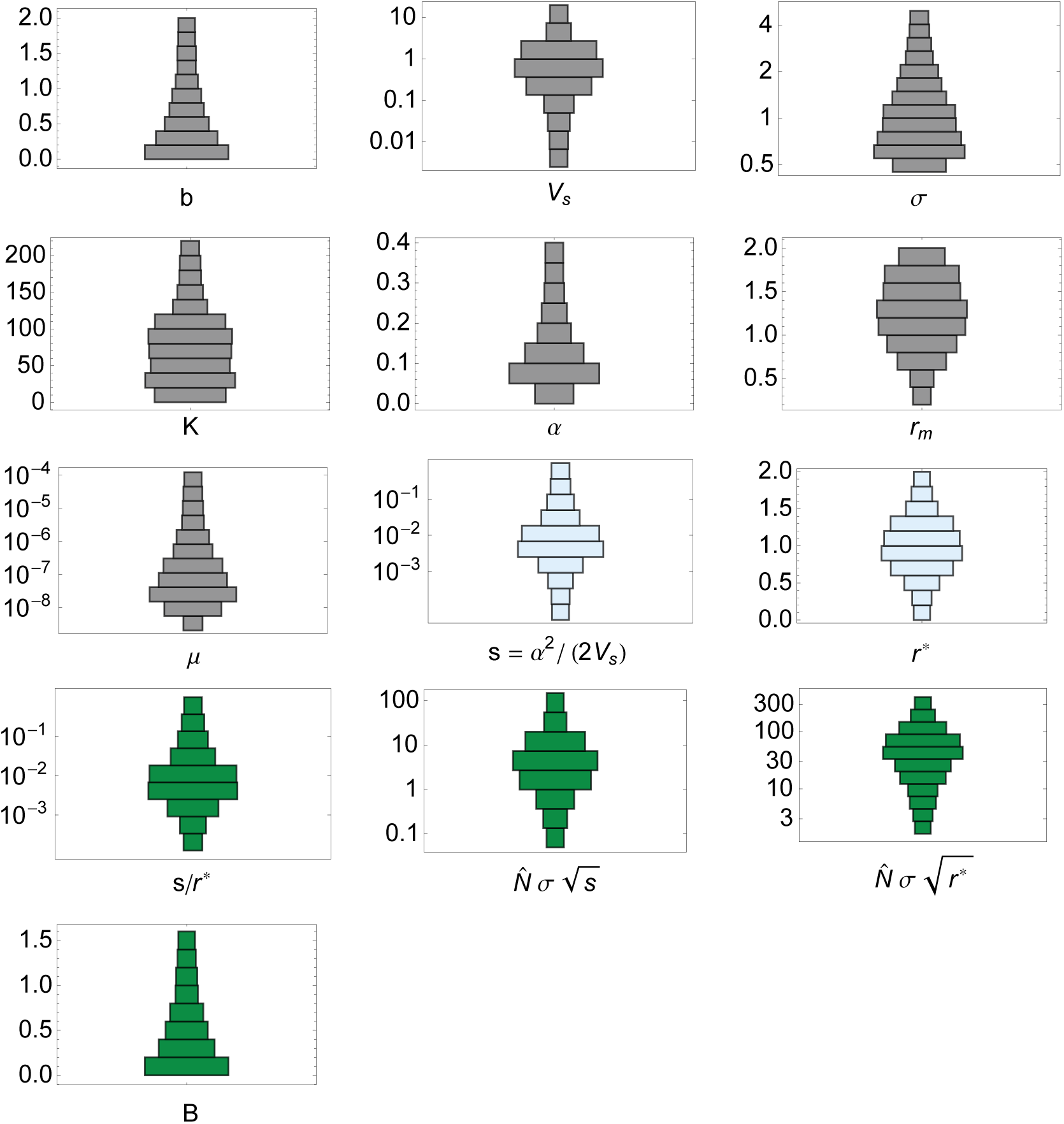
Distribution of parameters used in the “random set”, as described in Table 1. Grey: all unscaled parameters. Light blue: Composite parameters. Dark green: Scale-free parameters (both *s/r*^*^ and 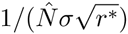 are given for a reference though one of them is always redundant; see Materials and Methods: Rescaling).

**Figure S2:**
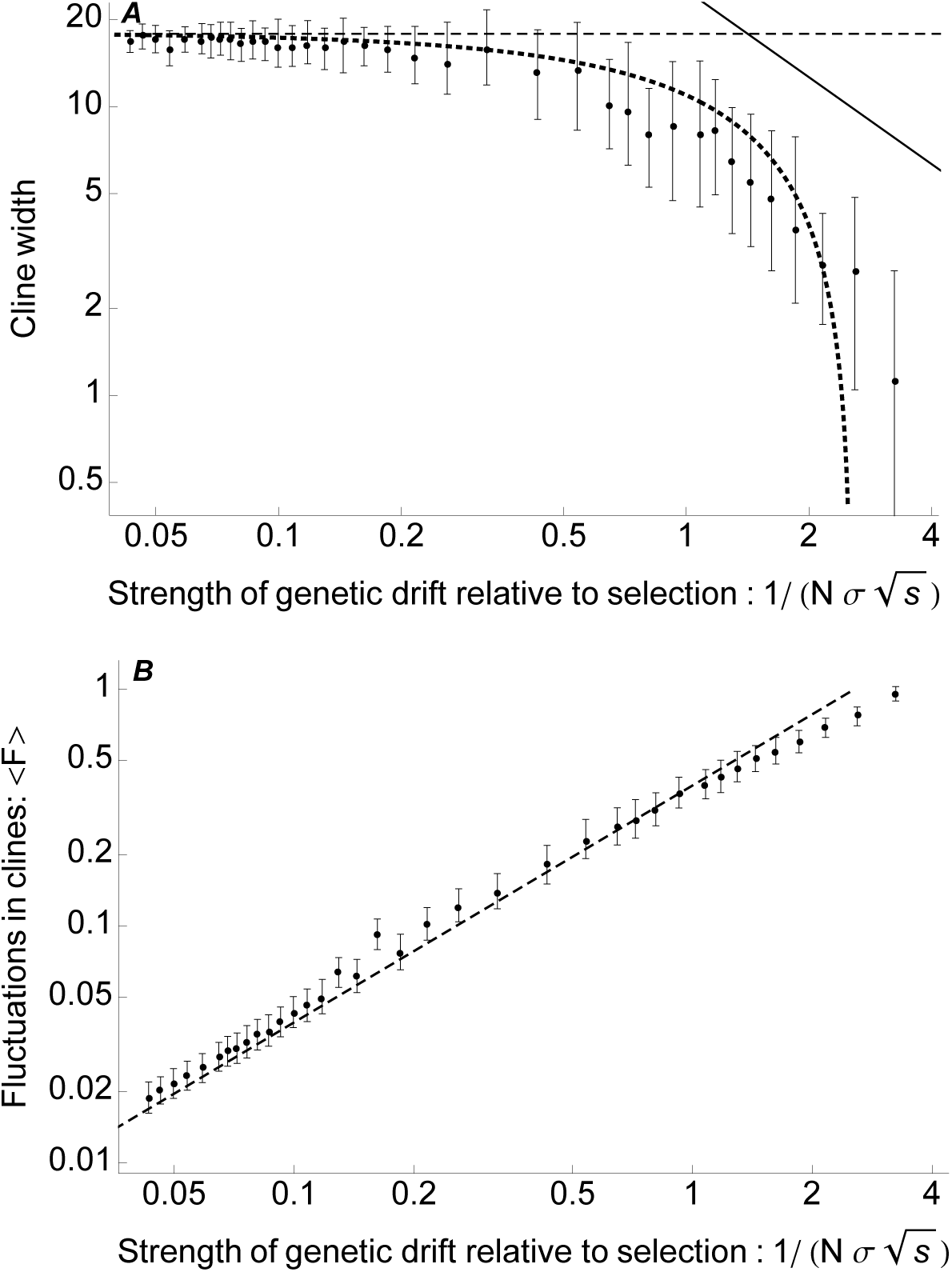
Cline width decreases as genetic drift gets stronger relative to selection. (A): Cline width, defined as a measure of total heterozygosity across space 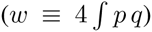, is approximated by 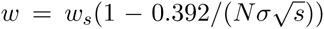 (thick dotted line; method from [3, supplementary text]). Dots give the median cline width across loci with their positive and negative standard deviations. The dashed horizontal line gives the deterministic cline width, 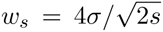 [8, 9]. As the genetic drift gets very strong relative to selection, the predictions diverges (bottom right): for very weak selection, the neutral limit 〈*w*_0_〉 → 4*σ*^2^*N* is approached (solid diagonal line; from [10]). Note that genetic variance decreases with the same factor as cline width. (B): The decrease in cline width can be understood from the rise in fluctuations of each cline – as fluctuations increase, allele frequencies fix locally and the cline steepens. Fluctuations in clines 〈*F*〉 rise approximately with 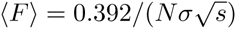, shown by a dashed line. 〈*F*〉 is the variance in allele frequencies scaled by the expected allele frequency, averaged across space, therefore ranging between 0 and 1: 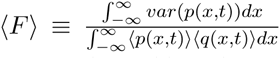 [3, p. 228]. Note that these formulae apply only to one-dimensional habitats: as the width of the second (selectively neutral) spatial dimension of the habitat increases, the predictions will start to differ because the effect of genetic drift on a cline depends only weakly on selection in two-dimensional habitats; see [11]). Parameters: *b* = 0.1, *σ*^2^ = 1/2, *V_s_* = 2, *r_m_* = 1.025, 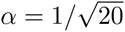, *μ* = 10^−4^, carrying capacity *K* increases from 4 to 260.

**Figure S3:**
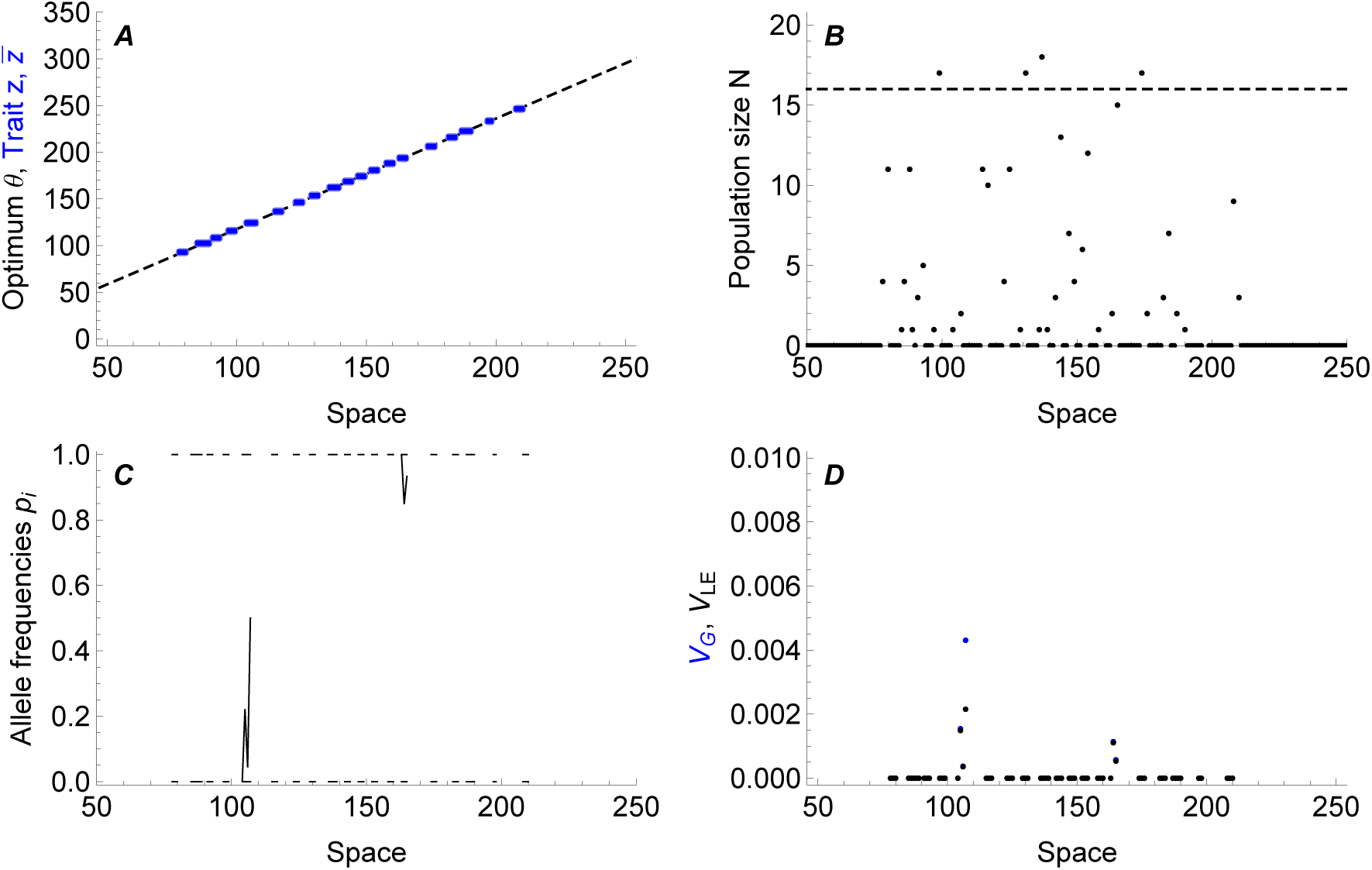
Species range can fragment when 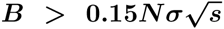 (Fig. 2), and additionally, 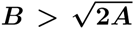 (Fig. 4). Exact conditions that always lead to range fragmentation were not determined. Fragmented populations are shown in open circles in Figs. 2 and 4. (A) Typically, the gradient in trait mean is zero within each fragment. (B) Populations are disjunct, and across the habitat, the population size is considerably smaller than predicted for “perfect” adaptation. (C) Typically, there is no clinal variation although transiently, a few clines are maintained. (D) Correspondingly, local genetic variance is mostly near zero. Parameters: *b* ≐ 1.18; *σ*^2^ ≐ 1, *V_s_* ≐ 0.44, *r_m_* = 1.97, *K* = 29.2, *μ* = 6 · 10^−8^, *α* = 0.0093, time = 5000 generations (shown at generation 4800). Note, that this fragmentation is not driven by edge effects that arise when population at the carrying capacity reaches the edge of the available habitat, where there is less maladaptive gene flow, which leads to local increase of density, followed by suppression of nearby populations towards the centre – and the effect propagates [12]. This effect is explained in [1]. Here, the simulations are set up such that the population never reaches the margins of the available habitat.

**Figure S4:**
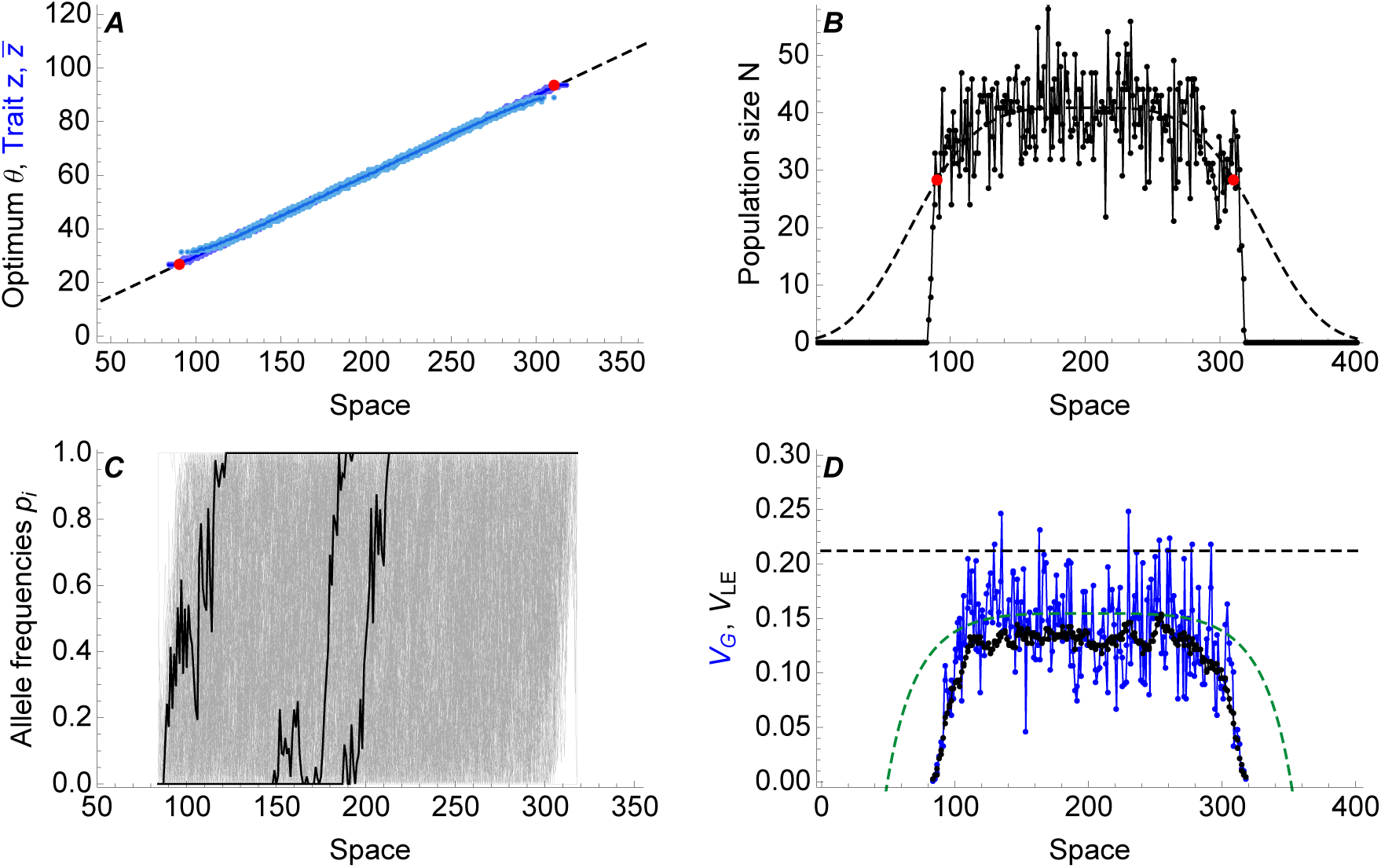
Non-uniform carrying capacity generates a stable range margin. (A) The optimum changes across the environment with a constant gradient *b* = 0.3 - the population starts well adapted at the more central part of the habitat (lighter blue). (B) The population density declines away from the centre - red dots give the predicted failure of the adaptation, based on 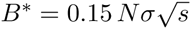. (C) Three representative clines are shown in black, other clines form the gray background (every tenth cline displayed). (D) Genetic variance is substantially lower than the deterministic prediction (black dashed line). The dashed green line gives the predicted 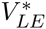 including the effect of genetic drift: 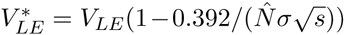(see Fig. S2). Parameters: *b* = 0.3, *σ*^2^ = 1/2, *V_s_* = 1, *r_m_* = 1.1, *μ* = 10^−7^. Population is shown after a stable range margin is reached at time = 100 000 generations (with the exception of the initial distribution of trait values, shown in light blue).

**Figure S5:**
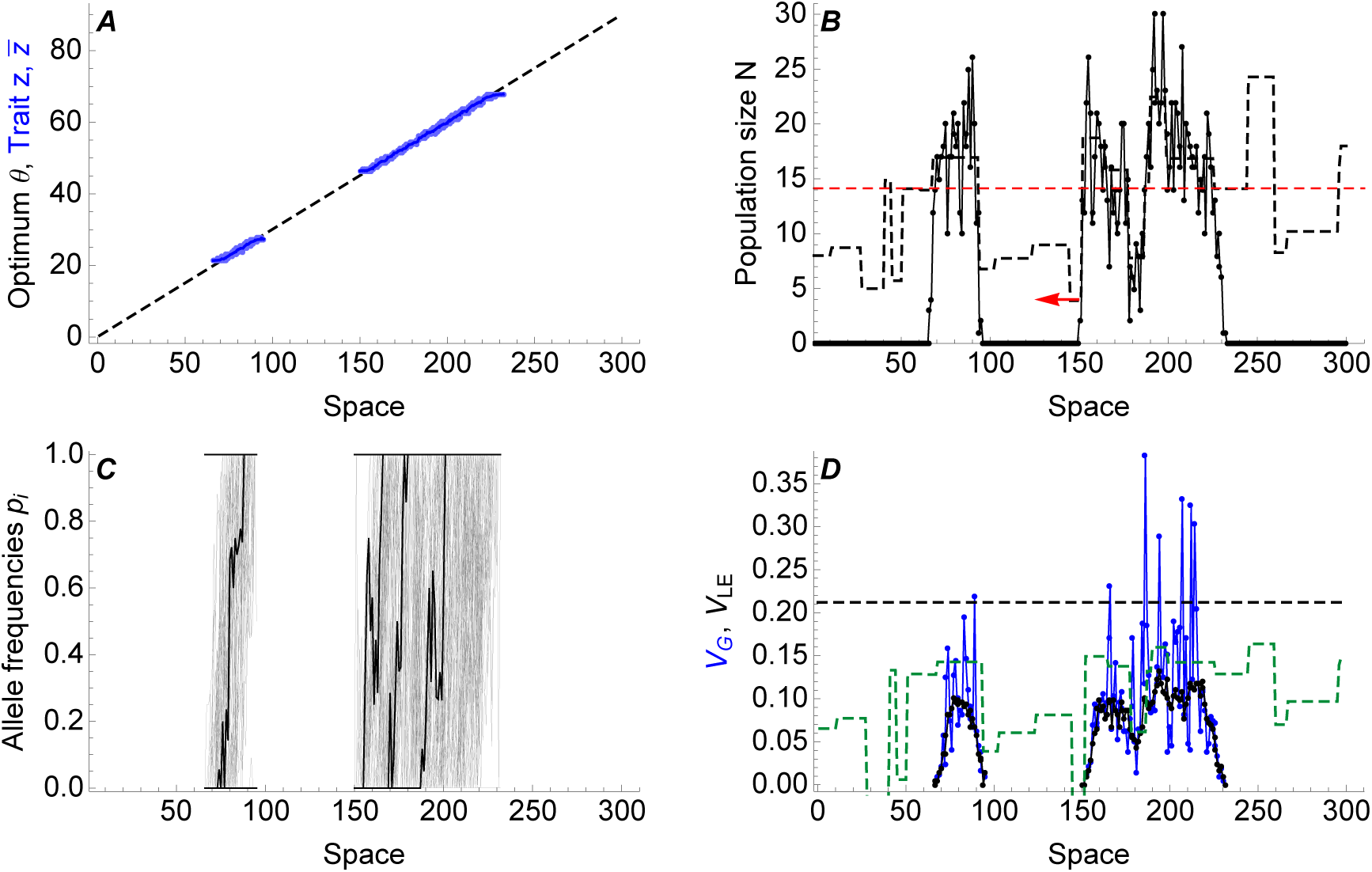
Species’ range is more robust within the already occupied habitat. A drop in the density to or below the predicted threshold 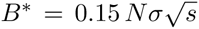(dashed red line in (B)) prevents range expansion. Yet, a considerably larger drop in the density is necessary before the range fragments within the occupied habitat (A, C). In this example, species’ range contracted from a local drop in the density right in the middle of the range (arrow in (B)), where locally, genetic drift depleted genetic variance to zero. Predicted segregating genetic variance 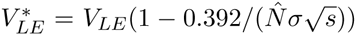 is given by green dashed line in (D). Note that segregating genetic variance 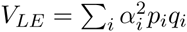 (connected black dots, D) changes over scales of 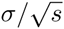, whereas population size (B) changes over much smaller scales *σ*. Species’ range is more robust to fluctuations within the already occupied habitat because migration from the neighbouring populations stabilizes the species’ range; the threshold (dashed red line) predicts when on a linear gradient, species’ range would collapse *from the margins*. Subplot descriptions and deterministic predictions (black dashed lines) are the same as in Fig. 1. Parameters: *b* = 0.3, *σ*^2^ = 1/2, *V_s_* = 1, 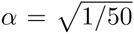, *r_m_* = 1.1, *μ* = 3 · 10^−7^, time = 40 000 generations. At the start of the simulation, population is adapted to the central half of the available habitat. Carrying capacity is generated from a Gaussian distribution centred around the threshold (*N*^*^, dashed red line) with a standard deviation of *N*^*^/3; the width of the patches is uniformly distributed between 1 and 30 demes.

**Figure S6:**
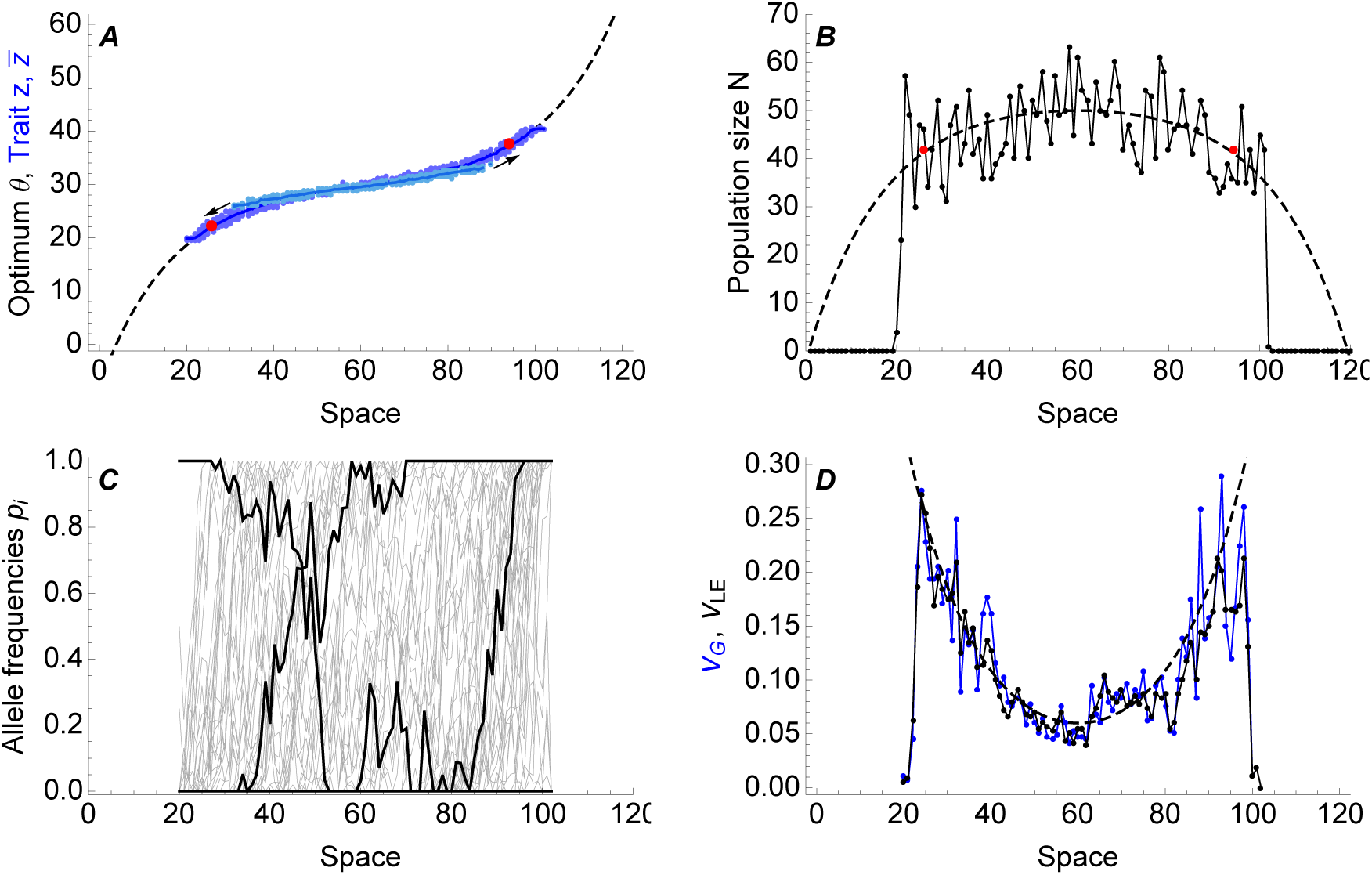
Sharp margin to a species’ range forms even when allelic effects *α_i_* are non-uniform. With exponentially distributed allelic effects *α_i_*, the expansion slows down after 40 000 generations (see Fig. S7 A), at the threshold predicted by mean selection per locus 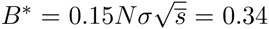 (red dots). As in this example the allelic effects are not bounded, over very long times, as rare alleles with large effect are recruited (see Fig. S7 B,C), the species’ range slowly stretches beyond the threshold *B*^*^. Parameters and subplot descriptions are the same as in Fig. 3 – with the exception that the allelic effects are exponentially distributed, with mean 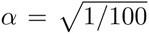. (C) Note that when allelic effects vary across loci, occasionally a cline may establish in a reverse direction, correcting a substitution with a large effect on the trait mean. (D) As genetic variance *V_G_* increases towards the margins, it evolves to match the ever steeper environmental gradient: 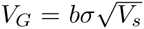, prediction shown in dashed line. Gradient in the central habitat is *b* = 0.12; *σ*^2^ = 1/2, *V_s_* = 1/2, *r_m_* = 1.06, *K* = 50, *μ* = 2 · 10^−7^, time = 100 000 generations (with the exception of the initial distribution of trait values, shown in light blue).

**Figure S7:**
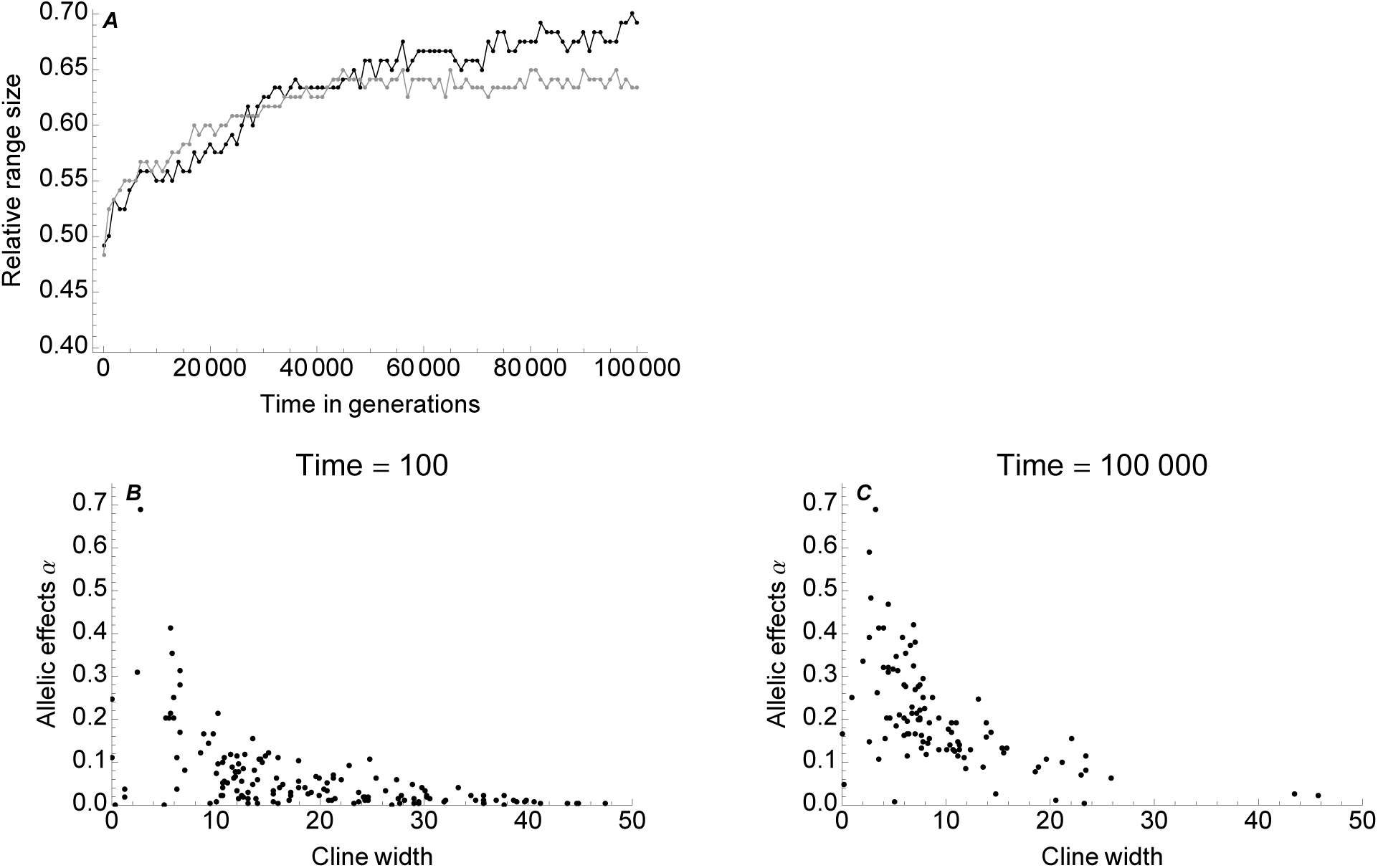
(A) Range expansion slows down near the threshold based on mean selection coefficient even when the allelic effects *α_i_* are non-uniform (black). Yet, over very long times, further alleles with large effects can be recruited as they are under stronger selection, and species’ range expands a little further. (Example from Fig. S6.) The extent of range expansion is only fully bounded by the substitution with the largest selection coefficient that can arise over a given time. For comparison, the rate of range expansion with equal allelic coefficients with 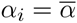 is given in gray (keeping all other parameters same, example from Fig. 3). (B, C) Over time, genetic drift degrades clines with small allelic effects *α*. As more alleles with larger effect *α* contribute to adaptation, clines become narrower and under stronger selection. Parameters as in Fig. S6. Note that the average selection coefficient is 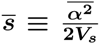 and in this example, *s* = *α*.

**Figure S8:**
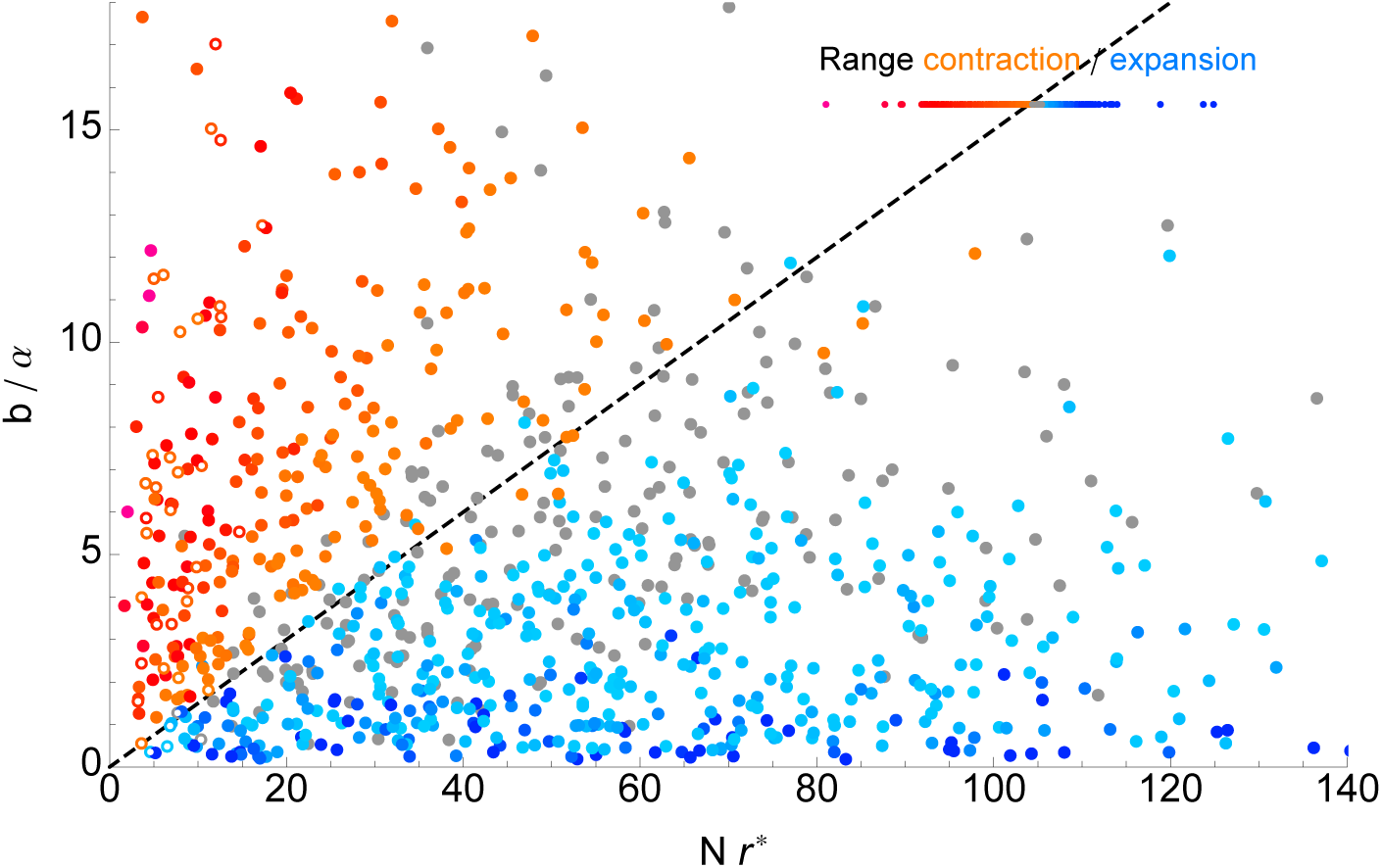
Alternatively, the threshold for range collapse can be expressed as *b/α* ≳ 0.15 *Nr*\*. The threshold holds well unless spacing between the clines, *α*/*b*, is smaller than about 1/10 - this reflects the limits of our simulation system rather than a biological boundary – the deme-spacing is fixed to Δ*x* ≡ 1. Data and their depiction are the same as in Fig. 2.

**Figure S9:**
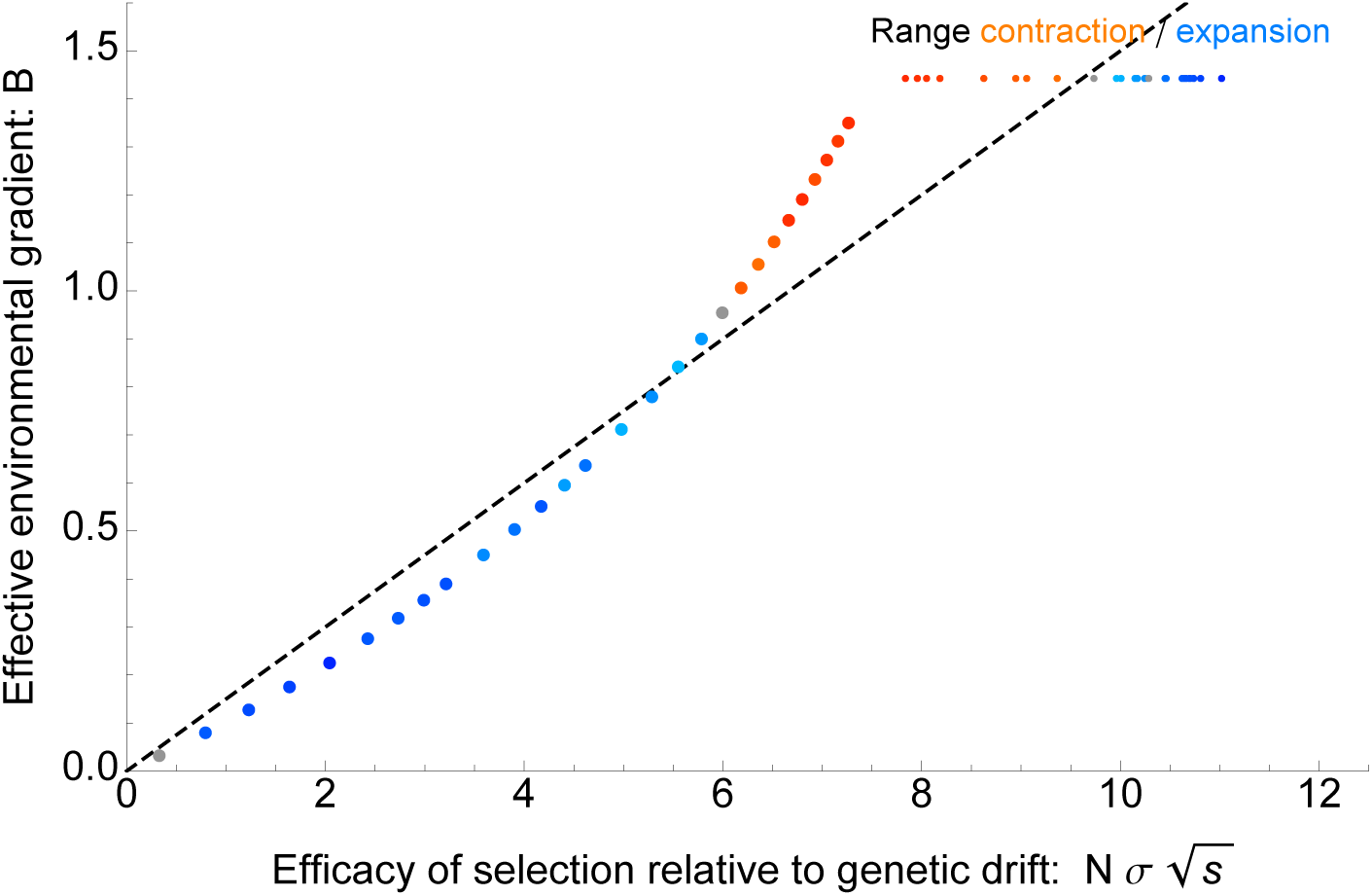
Adaptation may suddenly fail if dispersal is too large. The threshold for collapse of adaptation (dashed line, 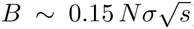) is weakly dependent on dispersal: to the first order, the effect cancels. Yet, for our model, the strength of density dependence *r*^*^ decreases with genetic variance (*r*^*^ = *V_G_/*(2*V_s_*)), which in turn increases with 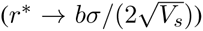. When 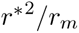 becomes smaller than 1, further increase in dispersal is detrimental as it brings the population closer to the predicted threshold. This is because both 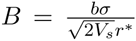 and 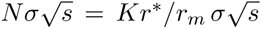 are both dependent on *r*^*^ – and the effects multiply. The scale for the colouring is adjusted (different to Fig. 2) so that the differences in the rate of range expansion (light to dark blue) and contraction (orange to red) are visible; gray dots again denote population that expanded less than one deme over 5000 generations. Parameters: *σ* = [0.1, 4.24], *b* = 0.45, *V_s_* = 1, *r*^*^ ≡ 1 hence *r_m_* = [1.02, 1.95]; *K* = 28, *μ* = 4 · 10^−7^, 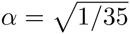, time = 5000 generations.

**Figure S10:**
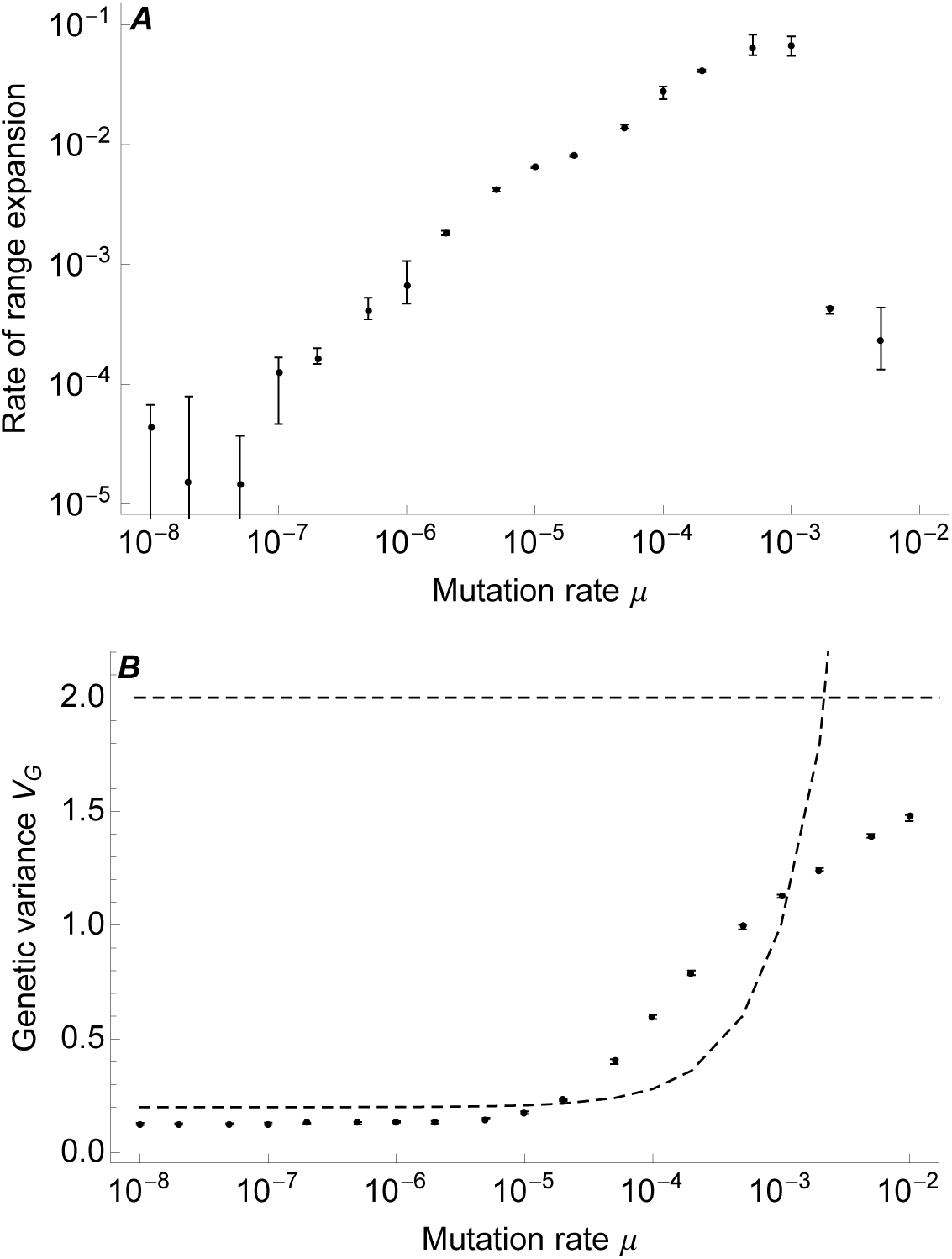
Effect of mutation rate on the rate of range expansion and local genetic variance. (A) Rate of range expansion increases about linearly with mutation rate *μ* per locus and generation (5 · 10^−8^ < *μ* < 10^−3^). When mutation becomes too high, then the rate of expansion first decelerates, and for yet higher mutation rates (*μ* ≳ 10^−2^), the population starts to collapse (c.f. [13]). Note that in general, mutation rate per locus and generation is expected to be lower than about 10^−4^ (see Supplementary Text 3). (B) Local genetic variance *V_G_* increases with mutation rate. For low to moderate mutation rates, genetic variance is maintained by mixing across the phenotypic gradient, 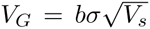. The dashed curve gives the prediction 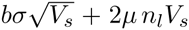, which assumes the components of genetic variance due to gene flow (first term) and mutation-selection balance (second term) combine additively, whilst all *n_l_* loci are at linkage equilibrium: the mismatch between the dashed curve and the realised genetic variance implies that with clinal variation, *μ* ≪ *s* is not a sufficient condition for the contribution of mutation to be negligible. The top dashed line gives the maximum variance possible, 1/4*n_l_ α*^2^ (where *α* is a phenotypic effect of a single substitution). With increasing variance, population density drops steadily (not shown): eventually, species’ range starts to contract and population collapses. Parameters: *b* = 0.4, *σ*^2^ = 1/2, *V_s_* = 1/2, *r_m_* = 1.06, *K* = 50, *α* = *s* = 0.01, 800 genes. Initial population spans over 100 demes, population evolves over 5000 generations. For both plots, error bars give the standard deviations.

